# Modeling robust and efficient coding in the mouse primary visual cortex using computational perturbations

**DOI:** 10.1101/2020.04.21.051268

**Authors:** Binghuang Cai, Yazan N. Billeh, Selmaan N. Chettih, Christopher D. Harvey, Christof Koch, Anton Arkhipov, Stefan Mihalas

## Abstract

Investigating how visual inputs are encoded in visual cortex is important for elucidating the roles of cell populations in circuit computations. We here use a recently developed, large-scale model of mouse primary visual cortex (V1) and perturb both single neurons as well as functional- and cell-type defined population of neurons to mimic equivalent optogenetic perturbations. First, perturbations were performed to study the functional roles of layer 2/3 excitatory neurons in inter-laminar interactions. We observed activity changes consistent with the canonical cortical model (Douglas and Martin 1991). Second, single neuron perturbations in layer 2/3 revealed a center-surround inhibition-dominated effect, consistent with recent experiments. Finally, perturbations of multiple excitatory layer 2/3 neurons during visual stimuli of varying contrasts indicated that the V1 model has both efficient and robust coding features. The circuit transitions from predominantly broad like-to-like inhibition at high contrasts to predominantly specific like-to-like excitation at low contrasts. These *in silico* results demonstrate how the circuit can shift from redundancy reduction to robust codes as a function of stimulus contrast.

## Introduction

The nervous system in general, and mammalian neocortex in particular, exhibit staggering complexity. Just in the visual cortex of the mouse, recent studies characterized ~100 transcriptomic (Tasic et al. 2016) and 46 morpho-electric cell types (Gouwens et al. 2019). However, the 6-layered structure is relatively similar across areas and species (Hill and Walsh 2005). How does this circuit represent the visual stimuli? Is the representation of visual input robust to noisy environments? Is the coding of the cortical circuit efficient enough for the representation of visual inputs? One approach to characterize cortical function is to consider the cortex as a shallow hierarchy of areas (J. A. Harris et al. 2019), with each area preforming a set of transformations from their inputs to their outputs that are a subset of possible transfer functions.

Since a comprehensive experimental characterization of the input/output transfer functions is out of reach, we seek to perform such a characterization using a comprehensive model of one cortical area. We have constructed such a model for the mouse primary visual cortex (area V1) at two levels of neuronal granularity, generalized leaky integrate-and-fire neurons (GLIF) and biophysically-detailed neurons with spatially extended dendritic trees (Billeh et al. 2020). The model is thoroughly constrained by experimental data in terms of distribution of cell types (Teeter et al. 2018; Gouwens et al. 2018), connectivity and visual inputs (Durand et al. 2016), and reproduces a variety of *in vitro* and *in vivo* observations of cellular activity under both two-photon calcium imaging as well as Neuropixels recordings (Siegle et al. 2019; de Vries et al. 2019).

The construction of the cortical model focused on reproducing *in vivo* activity. Here, we study how our computer model responds to perturbations, and we compare the responses to published experimental perturbations at the level of cell types (Olsen et al. 2012), single cells (Chettih and Harvey 2019) and functional populations (Marshel et al. 2019).

Experimental perturbations have been used to study functional interactions within populations *in vivo*. One of the questions asked is whether lateral interactions among excitatory neurons are dominated by like-to-like excitation or inhibition. Structurally, a salient feature of the observed connectivity is like-to-like in both probability (Gilbert and Wiesel 1989; Ko et al. 2011) and strength (Cossell et al. 2015) between L2/3 excitatory neurons. This has led to the proposal of a functional amplification role for these connections (K. D. Harris and Mrsic-Flogel 2013). However *in vivo*, predominantly a like-to-like inhibition has been observed (Vinje and Gallant 2000; Chettih and Harvey 2019).

We are interested in relating these perturbations with normative theories of cortical processing. One influential theory is that local circuits process the information efficiently (Barlow 1961; Attneave 1954). As beautifully reviewed by (Chalk, Marre, and Tkačik 2018), two regimes of efficient coding are described. For low noise, there is a need for redundancy reduction. One mechanism to implement it is functional like-to-like inhibition (Olshausen and Field 1996b; King, Zylberberg, and DeWeese 2013), which leads to the formation of a sparse code. However, at high noise, some redundancy needs to be preserved for optimal coding (Karklin and Simoncelli 2011; Doi and Lewicki 2014; Brinkman et al. 2016; Iyer and Mihalas 2017) which can be implemented with like-to-like excitation. Can the detailed cortical model match the observed perturbations, and can it balance redundancy reduction and robustness?

In this study, we used this newly-built V1 model employing the GLIF neuron (Teeter et al. 2018) representation to simulate experiments. It requires three orders of magnitude less computing time than the biophysically detailed version (Billeh et al. 2020; Gouwens et al. 2018), allowing us to more rapidly explore the space of inputs by conducting thousands of simulations, permitting exploration of the characteristics and coding mechanisms of V1 in an efficient way.

Starting from the responses to visual inputs, we investigate the effects of three types of optogenetic perturbations, simulated by direct de- or hyperpolarizing current injections. First, perturbations were performed at the cell type population level, mimicking cell type specific optogenetic perturbations to study the functional roles of excitatory layer 2/3 inter-laminar interactions. We provide access to comprehensive stimulations of perturbations of all cell types in the supplementary material. Second, single neuron perturbation simulations were conducted to explore how activity change of one neuron influences nearby neurons and network activity. The simulation results are generally consistent multiple features of the experimental observations (Chettih and Harvey 2019) however, specific higher order interactions differ. Finally, multiple-neuron perturbations were performed for co-tuned excitatory neurons of layer 2/3. While the neurons with similar tuning properties are affected by the perturbations, populations of excitatory neurons across tuning and retinotopic locations are barely affected. These changes are consistent with inhibition stabilization of the activity (Ozeki et al. 2009). The simulation results reveal a transition between a specific like-to-like excitation to a broad like-to-like inhibition when transitioning from low to high contrasts, and as a function of the size of the perturbation.

While it is desirable to have optimal behavior at both high and low noise levels, it is unclear how complicated the underlying structure must be to realize such a transition. We here demonstrate that our recently published biologically realistic model of mouse V1 (Billeh et al. 2020) precisely shows such a transition without requiring any alteration or parameter tuning. This shows the power of realistic models to generalize, and link to theoretical aspects outside of those for which they were trained.

The models were constructed and simulated using the Brain Modeling ToolKit (BMTK; https://alleninstitute.github.io/bmtk/) (Gratiy et al. 2018) interfaced with NEST (Peyser et al. 2017) and utilized the SONATA modeling format (https://github.com/AllenInstitute/sonata) (Dai et al. 2019). These tools and our simulation results are publicly available as a free resource for the community.

## Results

The GLIF V1 model, detailed in (Billeh et al. 2020), is visualized in Fig. 1A-C, and briefly outlined below. The model includes neurons in five layers, as layers 2 and 3 are combined as is standard for mouse cortex (Wang et al. 2020). Layer 1 has one (inhibitory) cell type, Htr3a, while all other layers are each populated by one excitatory and three inhibitory (Pvalb, Sst and Htr3a) cell types (Fig. 1A; 17 cell types in total). The network receives simulated visual input from the lateral geniculate nucleus (LGN), in addition to simulated background (BKG) from other cortical regions and experimentally imposed perturbation (PTB) that mimic the effect of optogenetic manipulations (Fig. 1C). The area of visual cortex that the model covers (Fig. 1A) contains a “core”, the portion of the model considered in all analyses here, a “periphery” that supplies extensive connections into the “core” to prevent boundary artifacts in the latter. The distribution of different neuron populations is visualized in Fig. 1B. All parameters are set as in (Billeh et al. 2020); single cell parameters were obtained from patch clamp measurements (Teeter et al. 2018); connectivity was constrained by literature and fit to reproduce background activity and evoked responses to drifting gratings, as described in (Billeh et al. 2020). No parameters were tuned to match known perturbation experiments for this study.

**Figure 1.**
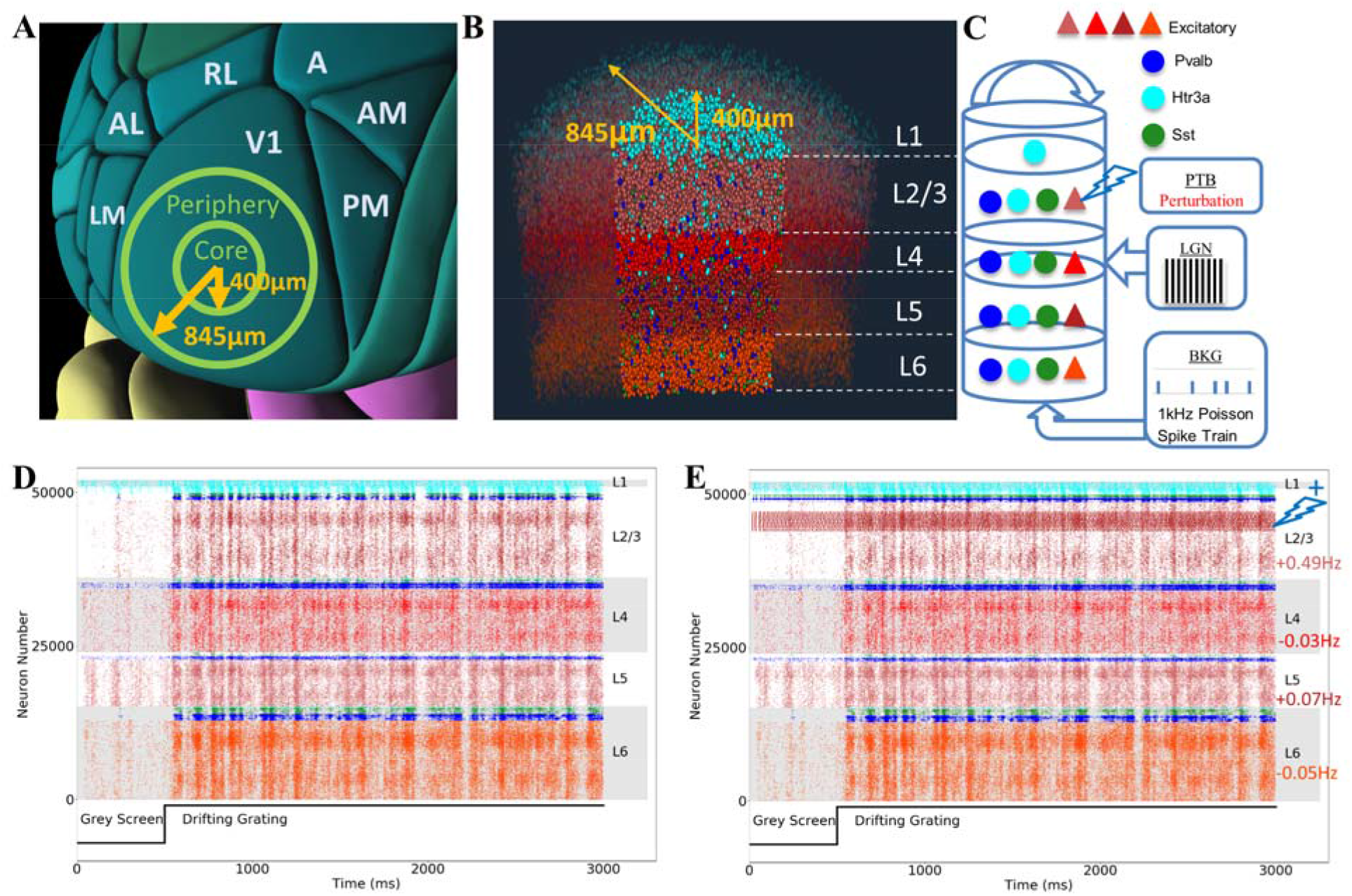
Characterization of the GLIF V1 model used. (**A**) Visualization of mouse posterior cortex illustrating the cortical surface area of V1 covered by the model (400 μm radius for the “core” within which neurons were analyzed and 845 μm radius with the surrounding annulus). (**B**) Visualization of half of the model to illustrate its composition. (**C**) Schematic diagram of the inputs from the lateral geniculate nucleus (LGN), background (BKG) and perturbation (PTB) with layers aligned to (B). Layer 1 contains a single inhibitory class of Htr3a. All other layers have an excitatory population and three inhibitory classes: Paravalbumin (Pvalb), Somatostatin (Sst), and Htr3a. (**C**) The input from the LGN projects to all layers in a cell-type specific manner, as constrained by experimental data (Ji et al. 2015). The model receives a 1kHz Poisson spike train background (BKG) input to simulate the collective influence from other areas of the brain. The perturbation current is injected by the PTB input to target cells. Inhibitory neuron types: Pvalb (blue), Sst (green) and Htr3a (cyan); colors are the same in each layer. Excitatory neurons are colored in different hues of red across layers 2/3, 4, 5, and 6. (**D**) Raster plot of a 3 s simulation of the model with LGN input as 0.5 s of grey screen, followed by 2.5 s of a drifting grating. Note the neuron numbers for every population are ordered based on the preferred their direction of motion. (**E**) Raster plot from a simulation where a subset of E2/3 neurons were activated (for the same stimulus as (D)). The perturbations applied to neurons that prefer motion 270°+/−45° and were situated within 100 μm from the center of the model. The injected current was 3 times rheobase of the target population. An activity stripe in the perturbed neurons in layer 2/3 is visible, with visual tuning being retained. The average Δ*f* for the entire E2/3 population during the grating period increased by 0.49 Hz, leading to a barely perceptible increase of 0.07 Hz in all E5 neurons and a tiny decrease of 0.03 Hz and 0.05 Hz respectively in all E4 and E6 populations. While there is a large increase in the rate of the stimulated neurons, and excitatory neurons at similar orientations and retinotopic locations, this causes an increased activation of inhibitory neurons, and a decrease in the rate of excitatory neurons at other retinotopic locations or orientations. These changes translate to a near balance at the level of the entire population, known as *inhibition stabilization* (Ozeki et al. 2009). The color code for neuron types in (B, D, E) is the same as in (C).

The majority of simulations are based on perturbations during visual stimulation by a drifting grating (with the following stimulus parameters, TF = 2Hz, SF = 0.04 cpd, contrast = 80%, orientation = 270°), from 0.5 s to 3 s, or during the presentation of a grey screen in the first half second. The stimulus conditions are varied while studying the effect of contrast on the perturbations.

### Cell Type Perturbations

We performed cell type specific optogenetic perturbations by injecting currents (either negative or positive) into the neurons of the targeted population (Fig. 1C). We conducted two types of perturbations: (i) complete silencing of an entire cell type population, and (ii) titrated activation of a subset of cells within a cell type (with E2/3 as an example). The results of the simulations are shown in Figs. 1E, 2, S1-S4.

**Figure 2.**
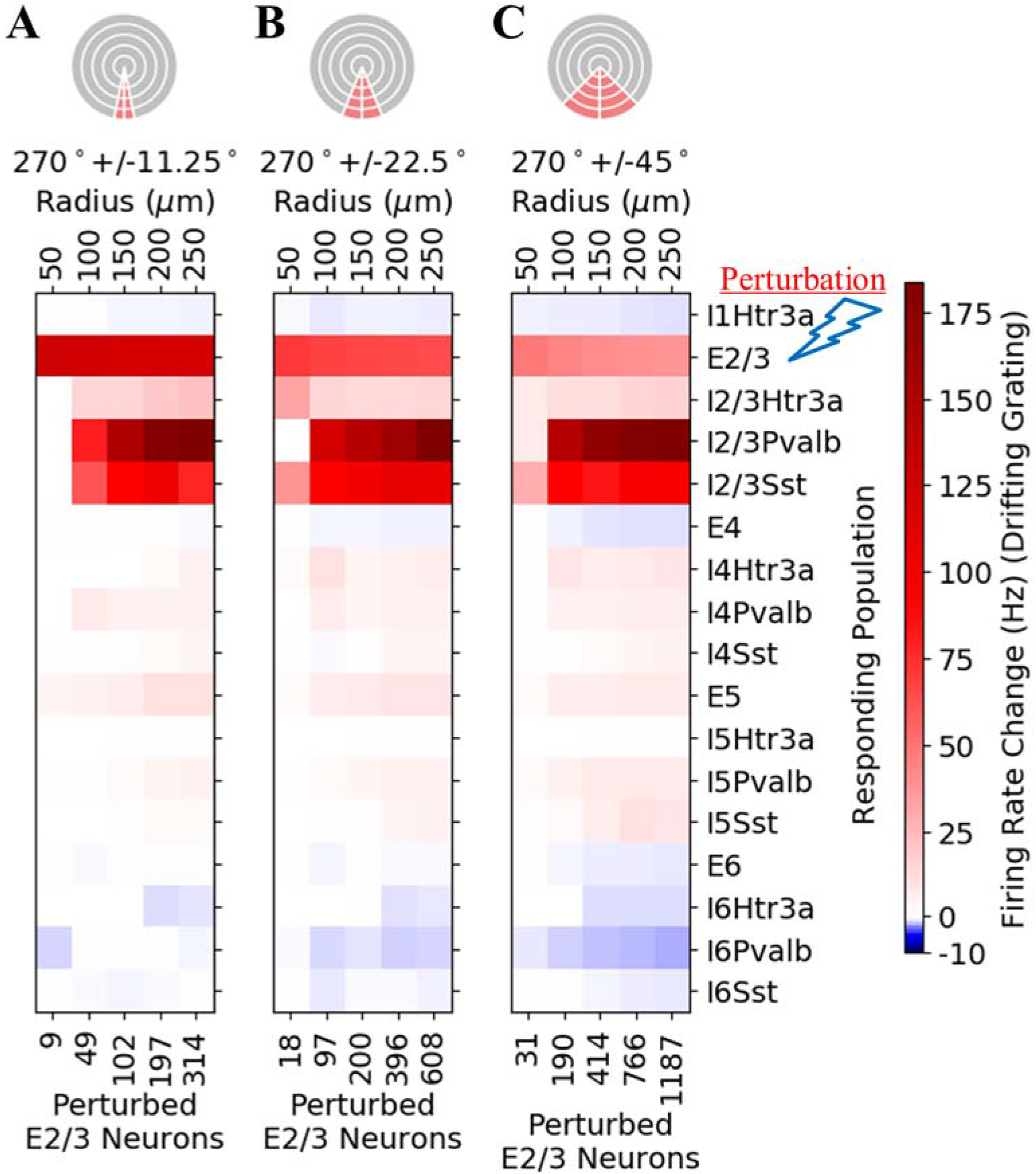
Functional population perturbations of excitatory neurons of layer 2/3 with the presentation of a drifting grating (TF = 2Hz, SF = 0.04 cpd, contrast = 80%, orientation = 270°). The injected current is three times each neuron’s rheobase. Perturbations were applied to neurons that prefer motion in the direction of (**A**) 270°+/−11.25°, (**B**) 270°+/−22.5°, (**C**) 270°+/−45°, within different radii (i.e., 50, 100, 150, 200 and 250 μm) from the center, as illustrated in the pie plot at the top. The activity changes (i.e., Δ*f*) are shown as heatmaps. The labels along the top x-axis of these heatmaps correspond to the radii of the relevant populations that are being perturbed with the number of neurons being perturbed at the bottom. The y-axis indicates the population for which Δ*f* is evaluated. Δ*f* is computed using only the neurons that have the same direction preference and radial positions as the perturbed population. The figure indicates successful imprinting by exciting the layer 2/3 population, leading to distinct patterns of activation. As expected, the effect of the perturbation becomes stronger as the number of perturbed neurons increases.

The main analysis metric we are using is firing rate change, 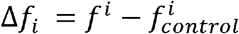, computed for every cell and then averaged over all cells within the analyzed selection of cells, where *f^i^* is the firing rate of neuron *i* during the perturbation, and 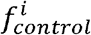 is the rate without the perturbation. Additional metrics are discussed in the supplementary material.

One experimental cell type perturbation study which we can compare simulation results against focused on layer 6 (Olsen et al. 2012). Positive current injections into layer 6 excitatory neurons (E6; Figs. S2A and S2B) show that these cells play a mildly inhibitory role on the upper layers via layer 6 Pvalb cells, which project to supragranular layers; correspondingly, suppressing layer 6 excitatory cells or layer 6 Pvalb cells results in disinhibition in upper layers, consistent with (Olsen et al. 2012).

### Functional Population Perturbation

Perturbation of a whole cell type may not reflect biologically relevant dynamics. With recent perturbations (Chettih and Harvey 2019; Packer et al. 2012; Yang et al. 2018; Peron et al. 2020; Carrillo-Reid et al. 2019) targeting smaller populations or even individual neurons, we investigated how perturbations of sub-populations may affect our V1 circuit. While we are not addressing the question of whether a percept emerges from the subjective perspective of an animal as a result of such targeted perturbations, we are investigating how the external “optogenetic” perturbation limited to neurons that share common response preference affects network dynamics.

We applied perturbations to groups of neurons that prefer the same or similar directions of motion of drifting grating: 270°+/−45°, 270°+/−22.5°, or 270°+/−11.25°. Furthermore, the perturbation was limited to neurons occupying the central portion of the model within radii of 50, 100, 150, 200 or 250 μm. Due to retinotopy, this selects for of neurons that share preferences for a small region of visual space. With an average cortical magnification of 70°/mm in the azimuth, these correspond to simulating 3.5 to 17.5° of visual field (Kalatsky and Stryker 2003; Schuett, Bonhoeffer, and Hübener 2002). All these perturbations were applied to a subset of E2/3 neurons. Ten trials of the drifting grating stimuli were run for each perturbation simulation, with results shown in Figs. 1E, 2, S3-S6.

It should be noted that while the effects of the perturbation on the firing rate of stimulated cells as well as the cells of the same orientation and same retinotopic location in other populations can be big (Fig. 2), due to inhibition of cells at different retinotopic locations or of dissimilar orientation, the effects at the level of the population are very small (Fig. 1E, Figs. S3-S5). The quantitative analysis of the inhibition of cells at different retinotopic locations and at different orientations for one stimulated cell is described in Fig. 3, and their dependence on the number of neurons activated and the contrast of the visual stimulus are described in Fig 4.

**Figure 3.**
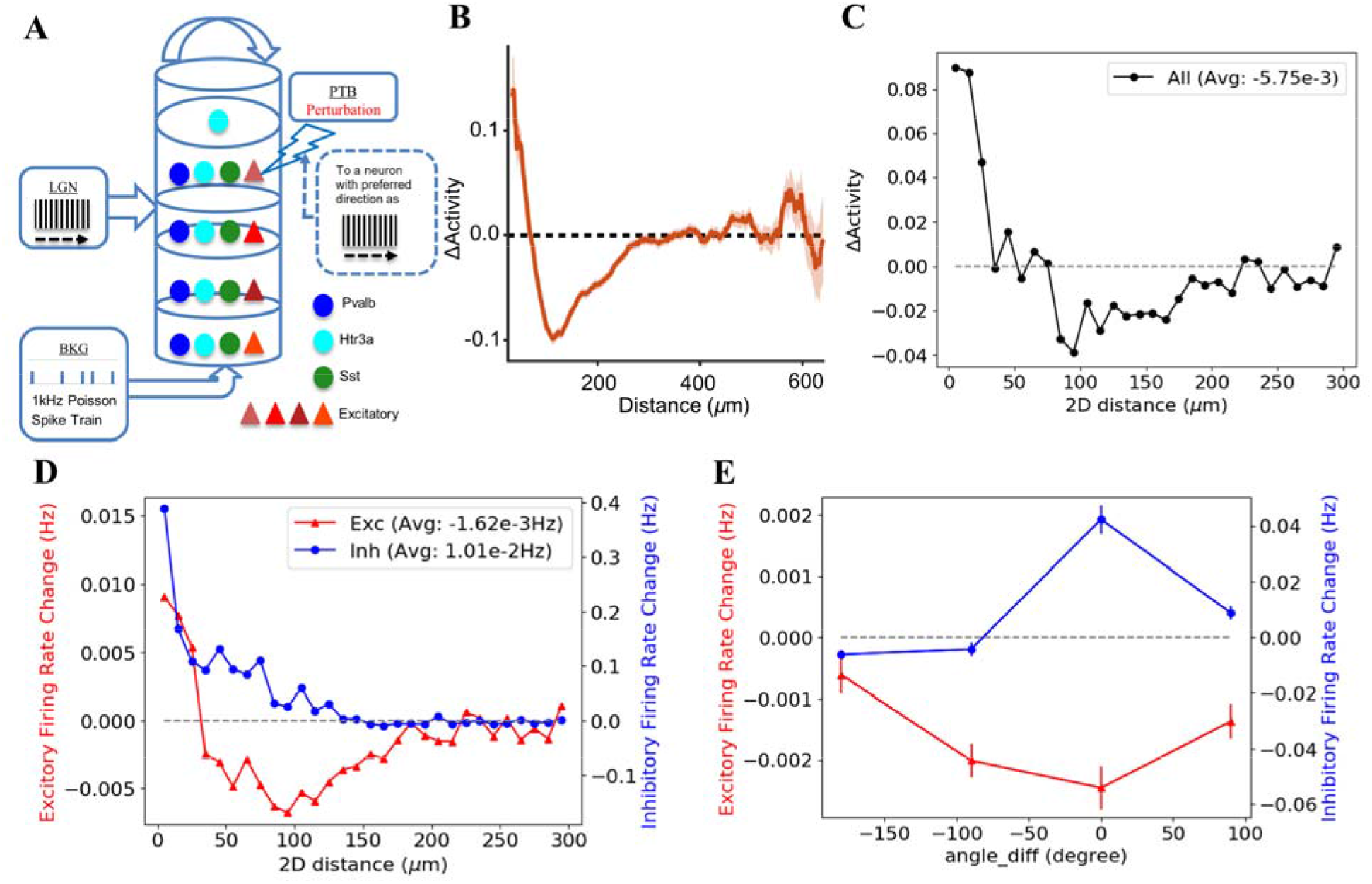
Using the V1 model to study effects of single-neuron perturbations on the population activity. (**A**) Perturbation of a layer 2/3 excitatory neuron, with the same preferred direction as the input stimulus. (**B**) Distance dependent the activity change from the experimental data (Chettih and Harvey 2019). (**C**) Combining the activity change of inhibitory and excitatory neurons using the metric from (Chettih and Harvey 2019) depending on the distance (in 10μm bins) from the single optogenetically stimulated neuron. (**D**) Change in firing rates depending on the distance (in 10μm bins) from the single optogenetically stimulated neuron. (**E**) Change in firing rates depending on the difference in preferred angle (in 90° bins) between the recorded and the single optogenetically stimulated neuron.

**Figure 4.**
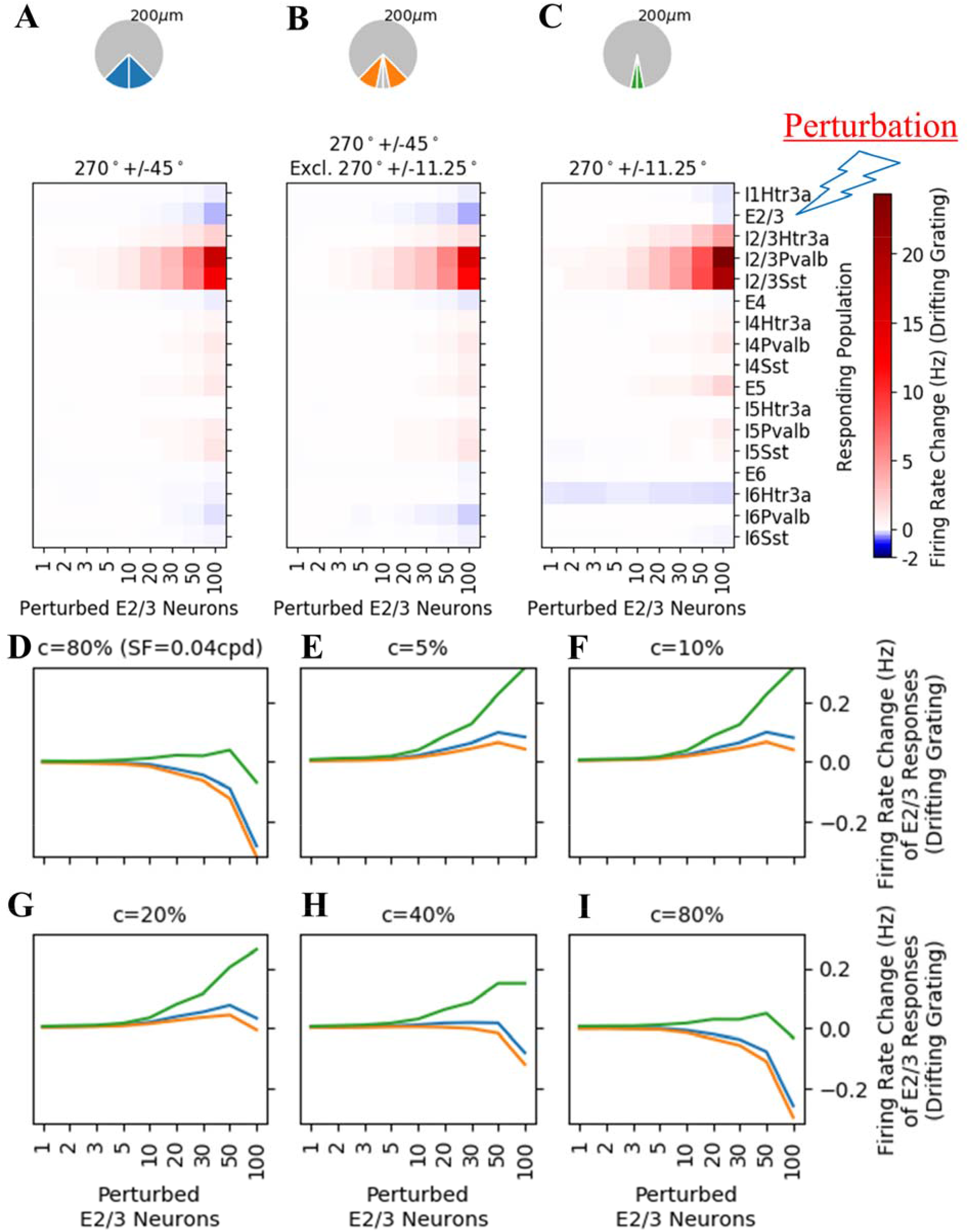
Perturbations of multiple excitatory neurons (from 1 neuron to 100) in layer 2/3 during drifting grating stimulation. The perturbed neurons were selected such that their preferred direction closely matched the direction of the drifting grating: 270°+/−11.25° and within 200 μm radius. (**A-C**) Analysis of neurons within a directional cone of 270°+/−45°, (**B**) 270°+/−45° but excluding 270°+/−11.25°, or (**C**) 270°+/−11.25°. The firing rate changes are shown as heatmaps. Labels along the y axis indicate the population for which the firing rate change is computed. For E2/3, stimulated neurons are excluded. E5 neurons were activated while the ones on other layers were suppressed. Inhibitory neurons were activated except those in layer 6. These effects increase as the number of perturbed neurons increases. (**D**) Activity changes of E2/3 neurons within different analyzed ranges. Closely aligned E2/3 neurons (within 11.25°) increase their activity for up to 50 stimulated neurons but decrease their activity thereafter. Similarly tuned E2/3 neurons (within 45°) show decreasing activities. (**E**-**I)** Activity changes of E2/3 neurons within different analyzed ranges for drifting grating with 0.02 cpd spatial frequency for five different contrasts (5%, 10%, 20%, 40% and 80%). At lower contrasts, closely-aligned E2/3 neurons increase their activity while similarly tuned neurons show a non-monotonic dependence on the number of neurons stimulated, with a maximum around 50 neurons.

As an illustrative example, the raster plot of one of the perturbations applied to E2/3 neurons that prefer motion in three directions (270°+/−11.25°, 270°+/−22.5° and 270°+/−45° within 100μm from the center of the model) is shown in Fig. S3. We can see stronger activity for the neurons being perturbed in Figs. 1E and S3 for E2/3 population. The more neurons perturbed, the bigger the band of elevated activity.

Fig. 2 shows the overall activity changes of these local perturbations. The figure illustrates Δ*f* for populations of neurons that share the direction preference and retinotopy with the perturbed neurons. Activating subsets of E2/3 cells leads to increased activity in co-tuned L5 neurons. This activity increases with the number of neurons stimulated up to ~50-100 neurons, saturating subsequently. This number is similar to the observed neuron number required for perception (Marshel et al. 2019). The stimulation also causes a massive increase in activity of the co-tuned inhibitory neurons in L2/3, and a general suppression of activity in L4 and L6. These patterns are similar to the expectations from the canonical cortical model (Douglas and Martin 1991). Also, of interest is the effect of these finely tuned perturbations on the entire cell types population (not just the co-tuned portion of it), shown in Fig. S4. Across orientations, the effect is mostly inhibitory and much weaker than the one in Fig. 2. We observe the same effect in Figs. S5 and S6 when using Optogenetic Modulation Index (OMI) rather than Δ*f*. This is expected in a circuit in which the recurrent inhibition stabilizes overall network activity (Ozeki et al. 2009).

### Single Neuron Perturbation

To compare results with a recent experiment (Chettih and Harvey 2019) that characterized the effects of single-neuron perturbations on the activity of nearby layer 2/3 neurons, we likewise activated individual neurons. Positive currents were injected to a target E2/3 neuron close to the center of the V1 model. Fifty sets of perturbation simulations, each done for a different target neuron, were conducted, with 10 trials each. The results are shown in Figs. 3, S7-9.

First, we perturbed an E2/3 neuron with the same preferred direction as the input stimulus (Fig. 3A). Of course, in our simulations we can measure small effects very precisely by fixing the seed of the random processes to be the same as in the unperturbed simulations. We analyze the simulation results using the same metric as in (Chettih and Harvey 2019), i.e., ΔActivity, defined as the firing rate changes divided by the standard deviation of such changes. This measures the change to the network relative to the trial-by-trial variability (e.g. a measure of 1 would mean that perturbing a neuron would create a change equal to the standard deviation of the trial-by-trial variability). To more directly compare with the experimental data, the values for ΔActivity are averaged for all neighboring neurons, no matter whether excitatory or inhibitory. Averaging over spatial positions and orientation tuning, we observe that, following the stimulation of one E2/3 neuron, there is a very modest decrease (−5.75×10^−3^) in the average relative activity of other L2/3 neurons. This perturbation is on the same order of magnitude as the experimentally observed one (−2×10^−3^). We can observe the trend of activity change with distance: nearby neurons (<50μm) were activated, while neurons further away (50-200μm) were suppressed, and the effect fading away with distance (Fig. 3C), showing similar spatial interaction terms to those experimentally observed (Chettih and Harvey 2019) also shown in Fig. 3B.

To delve into cell type specific effects, we analyze the results by separating excitatory and inhibitory neurons. We also use the firing rate change as a metric as it is easier to develop an intuitive understanding for the magnitude of the changes. Excitatory neurons follow a similar spatial dependency as the average, with nearby neurons being excited and the near surround inhibited (Fig. 3D). Inhibitory neurons follow a different pattern, lacking the near surround inhibition. As we averaged the effect over all orientation/direction preferences for excitatory neurons, distance-dependency is not dependent on the preferred angle of the stimulated neuron (Fig S7B). Since experimentally there is a potential bias towards recording excitatory neurons, we performed a more in depth analysis of how potentially different samplings of the different cell types can affect this result (Fig. S8). We observe that a weighted sum biased for excitatory neurons (Fig. S8) shows an even better fit to the experimental data than the average metric.

While the average changes in excitatory neuron activity seem small (−1.6×10^−3^ Hz), they are averaged over 3176 neurons. On a per neuron basis, stimulating one E2/3 neuron causes ~120 additional spikes (over 2.5 s), but leads to a *decrease* of 45 spikes discharged by other excitatory neurons throughout the column up to 300μm horizontally.

Beyond the dependence on relative spatial location (which in our model maps directly to retinotopy), the interactions are dependent on the relative functional similarity in terms of orientation/direction preference. When the visual stimulus is similar to the preferred stimulus of the stimulated cell, inhibition predominantly acts on similarly tuned neurons (Fig. 3E). The suppression of the similarly tuned neurons is larger than a normalization would predict: the relative decrease for similarly tuned neurons is ~3 times larger than the relative decrease for neurons of opposite direction.

We found it impressive that a model which did not have any parameters tuned for this computation was able to replicate these experimental results. However, no model is perfect. We found significant differences between experimental observations and the model results when looking at the interaction between the recorded and stimulated neurons when the visual stimulus, stimulated neuron and recorded neurons each had different preferred directions of motion (Figs. S8 and S9) which is discussed in the supplementary material.

### Multiple Neuron Perturbation

Further perturbations were done for multiple E2/3 neurons to study their effect on the surrounding neurons and the preferred angle of motion dependence of surrounding neurons. This is different from the functional population perturbation where we stimulated all neurons of a particular type and with a particular tuning. Positive currents were injected into multiple target E2/3 neurons (from 1 neuron to 100) around the center of the model (within a radius of 200 μm from the center), preferring approximately the same direction as the stimulus input (270° within a +/−11.25° range). Ten sets of perturbation simulations, each done for a different number of target neurons, were conducted, with 10 trials for each set (Fig. 4, S10).

We analyze the effect of these activations on three populations of neighboring cells (Fig. 4A-C). Neurons which prefer 270°+/−11.25° are very closely aligned in orientations to the perturbed neuron. Neurons preferring 270°+/−45° but excluding 270°+/−11.25° are similar but not closely aligned, correspond to a functional near surround. Neurons preferring 270°+/−45° are similar in tuning and represent the sum of these populations. In all these cases E5 neurons were activated while excitatory neurons in other layers were suppressed. This is consistent with the results shown in the 200 μm column in Fig. 2A, while being smaller in magnitude as fewer E2/3 neurons were perturbed. We also observe a general activation of inhibitory neurons except those in layer 6 (similarly to the results in Fig. 2). The effect increases as the number of perturbed neurons increases.

For E2/3 neurons, we observe a non-monotonic dependence (Fig. 4D). While neurons with similar tuning are suppressed irrespective of the number of perturbed neurons, closely aligned neurons are activated for a small number of perturbed neurons but show suppression if more than >50 neurons are triggered. This is caused by a “center-surround” organization in functional space (Fig S10), biased towards suppression. However, the effects of multiple stimulated neurons are not additive. An additional shift towards suppression is observed, with a transition towards suppressing everything else in the population when the number of stimulated neurons exceeds 50.

As a control, we analyzed the dependence of the response to perturbation on the spatial frequency of the stimulus. As the radius of the analyzed volume for this perturbation is 200 μm, given the average retinotopic magnification, this corresponds to 6° of visual field. With a 0.04 cpd spatial frequency of the stimulus, the maximal phase difference from the center corresponds to a 90° phase difference. We analyzed the response to a 0.02 cpd spatial frequency, in which the radius of the analyzed volume corresponds to 45° phase difference (Fig. 4E), and we observed practically the same response to the perturbations.

The pattern of functional lateral interactions significantly changes with the contrast of the visual stimulus. Here we used a drifting grating with spatial frequency of 0.02cpd with 5, 10, 20, 40 and 80% contrast (Figs. 4E-4I, Fig S10). At high contrasts, inhibitory interactions are prevalent. Similarly tuned E2/3 neurons (within 45°) monotonically decrease their activity as a function of neurons stimulated, while closely-aligned (within 11.25°) neurons had a non-monotonic dependence, with maximal activation for 50 stimulated neurons. At low contrasts, excitatory interactions are predominant. Closely-aligned neurons monotonically increase their activity as a function of neurons stimulated while similarly tuned neurons have a non-monotonic dependence with a maximum activation also for 50 neurons stimulated. The model shows a center-surround interaction in functional space (Fig S10).

This makes answering the question of whether we observe functional like-to-like excitation or inhibition in the model complicated. It depends on the contrast of the stimulus, the number of neurons stimulated and how strict one is in the definition of “like”. In general, at low contrasts we mainly observe specific like-to-like excitation and at high contrast a broad like-to-like inhibition.

## Discussion

In this study, we simulated optogenetic experiments using a previously constructed (Billeh et al 2020), anatomically and biophysically constrained model of mouse primary visual cortex to understand cortical processing. All parameters in the model, consisting of about 250,000 GLIF model neurons of 17 different cell types distributed from L1 to L6 (Fig. 1), were fixed as in Billeh et al (2020). We then activated small number of neurons, ranging from one to hundreds, and entire cell-type populations, to mimic optogenetic excitatory perturbations. The model is useful as a tool to quickly test outputs of potential experiments and thus help with experimental design.

As a direct effect of the perturbations of single neurons, we found that exciting single E2/3 neurons gives rise, on average, to a reduction in activity (Fig. 3). Such an effect increases with the size of the perturbation (Fig. S10). An inhibitory first-order effect with perturbation size is needed for inhibition stabilization of the global activity (Ozeki et al. 2009). Spatially, a center surround effect is observed. Regarding the dependence on the orientation, for single neuron perturbations we observed a broad inhibition which more strongly affects the more strongly responsive neurons. These findings are consistent with experimental data (Chettih and Harvey 2019). However, higher order interactions, in which the visual input, stimulated neuron and recorded neuron have different properties, are different in the model from experiments. One potential explanation for the difference is the complexity of the tuning of the neurons. Both in the experiment and the model, there is a “Mexican hat” interaction in functional space. I.e. when stimulating one neuron there is an excitation to closely-aligned neurons, and inhibition for similar neurons. However, in the experiments the number of closely aligned neurons is much smaller than in simulation, such that even when stimulating one neuron generally like-to-like inhibition is observed, even at lower contrasts. In simulations, to observe like-to-like inhibition we need to stimulate more neurons and provide a higher contrast stimulus than needed in the experiments.

We focused our analysis on the observed functional like-to-like inhibition when perturbing single neurons, as it is considered to be a hallmark of redundancy reduction (Olshausen and Field 1996b; King, Zylberberg, and DeWeese 2013). However, in the simulations it is not universal. It should be noted that no two neurons in the simulation have exactly the same tuning: they are randomly drawn from continuum distributions in retinotopic and orientation tuning. Anatomically, neurons have both like-to-like excitatory connections as well as like-to-like inhibition. Which of these anatomical connections ends up being dominant depends on the particular difference in tuning and is dependent on the contrast of the visual input. Generally, we observed more like-to-like excitation at low contrasts (Fig 4), congruent with models of recurrent amplification (K. D. Harris and Mrsic-Flogel 2013) and more like-to-like inhibition at high contrasts (Olshausen and Field 1996b; King, Zylberberg, and DeWeese 2013). These observations are consistent with a robust and efficient code with an emphasis on efficiency at high contrasts and robustness at low contrasts (Karklin and Simoncelli 2011; Doi and Lewicki 2014; Brinkman et al. 2016).

Beyond the general transition from more excitation at low contrast to inhibition at high contrast, the functional interactions are also dependent on how strict we are in the definition of similarity in tuning and the number of neurons stimulated. For closely-aligned neurons (<12.5 degrees difference) at high contrast (80%), and for similar neurons (<45 degrees difference) at low contrast (<20%) we observed a non-monotonic dependence of the perturbation effect on the number of neurons stimulated. In both cases the largest perturbation is observed when ~50 neurons are stimulated. This might form a critical mass of neurons needed to change the network state.

While the model incorporates a large amount of diverse neurobiological data, the model also has a large number of unconstrained parameters. We have not systematically explored the parameter space, but we provide a series of perturbations to all the cell types in the model as a battery of tests (Fig. S1). These tests, when/if the experiments become available, can be used by the community to further evaluate the predictive power of this model.

We are fully cognizant that no model is perfect. We view this work as a further refinement of a long line of existing models stretching back more than a century, combining models of cellular excitability (Lapicque 1907; Hodgkin and Huxley 1952) with models of sensory system processing (Fukushima 1980).

As a general conclusion of the study, as we have not tuned any new parameters for this circuit which we built with careful attention to local circuit properties and different single cell nonlinearities (Billeh et al 2020), the fact that we observed a transition from a broadly tuned like-to-like inhibition dominated circuit at high contrast to specific like-to-like excitation dominated interactions at low contrast, enables us posit that the local circuit transitions from redundancy reduction when signal-to-noise of the input is high, to robust code in the low signal-to-noise regime, as predicted to be needed by theories of efficient coding (Karklin and Simoncelli 2011; Doi and Lewicki 2014; Brinkman et al. 2016).

## Methods

The methodology of the V1 model building and perturbation simulation design is discussed in this section.

### Model Structure and Building

We briefly summarize here the V1 model structure (Billeh et al. 2020). The full V1 model described a cylinder of cortical tissue with a radius of 845 μm. The model was built with 230,924 neurons with 51,978 core neurons within the 400μm radius from the center of the V1 which are the main subject of analysis. The model spans all layers of V1 and has 17 different cell types (shown in Table 1). The diagram of the network is shown in Fig. 1C, together with structure illustrations shown in Figs. 1A and 1B. We used the Generalized Leaky Integrate and Fire (GLIF) version of the model as it is significantly faster to simulate.

**Table 1.** Cell Type Populations Included in the GLIF V1 model

|  |  |  |  |  |  |  |  |  |
| --- | --- | --- | --- | --- | --- | --- | --- | --- |
| E2/3 | E4 | E5 | E6 | I1Htr3a | I2/3Htr3a | I2/3Pvalb | I2/3Sst |  |
| I4Htr3a | I4Pvalb | I4Sst | I5Htr3a | I5Pvalb | I5Sst | I6Htr3a | I6Pvalb | I6Sst |

The model receives input from an LGN network of 17,400 nodes and a background source. There are totally 3,506,880 connections from LGN to the model, in which there are 786,405 connections to the core nodes at the center from LGN. The neurons in the V1 model are also recurrently connected with each other. The total number of recurrent connections within the V1 model is 70,139,111, and the background node has 230,924 connections to all the nodes in the V1 model. Note that, each edge has a different number of synapses.

#### GLIF Node Model

A class of point neuron models, i.e., GLIF (Generalized Leaky Integrate-and-Fire) models, were recently used to fit the responses of a large number of neurons of different cells types in the mouse visual cortex (Teeter et al. 2018). In this study, we used Level 3 GLIF (GLIF3, or termed LIF-ASC, i.e., leaky integrate-and-fire with after-spike currents) model, which was formulated as

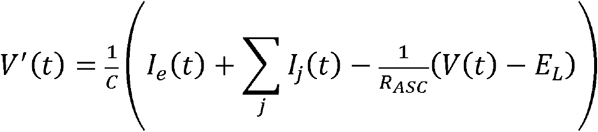

where *V*(*t*) is the membrane potential and *I*_*e*_ is the external injection current. When *V*(*t*) > *θ*_∞_ (with *θ*_∞_ as the instantaneous threshold), *V*(*t*) is reset by the rule of

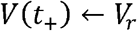

with the resetting potential *V_r_* = *E_L_*. The GLIF3 model considers the ion currents *I_j_*(*t*) activated by a spike, which is formulated as

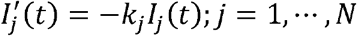

The update rule, which applies if *V*(*t*) > *θ*_∞_, is given by

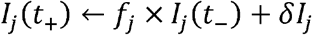

where multiplicative constant 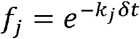. In this study, we use two sets of after spike current parameters in the GLIF3 model (i.e., *N* = 2). For detailed description of GLIF3 models and other four GLIF models, please refer to (Teeter et al. 2018).

The parameters for the GLIF models were trained based on the intracellular electrophysiological data. The V1 model was built based on 111 cell models for the 17 populations shown in Table 1, with parameters for each of these 111 cell models available through Allen Cell Types Database (ACTD) (Allen Institute for Brain Science 2017).

#### Synapse Mechanism

In this study, the NEST Simulator (Peyser et al. 2017) implementation of GLIF3 model was used through BMTK (Gratiy et al. 2018). Linear exact solution and interpolated spike time were set in the GLIF3 model for the V1 model simulation. Postsynaptic current-based synaptic ports were used for all the GLIF3 models to take inputs from LGN, background, recurrent connections within the V1 model. An alpha shape function was defined for the postsynaptic current (PSC) as

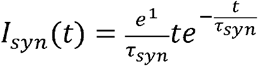

where *τ_syn_* is the synaptic port time constant. This function was normalized such that a postsynaptic current with synapse weight one has an amplitude of 1.0 pA.

The synaptic port time constants *τ_syn_* for different types of connections are defined list as follows. These time constant values were estimated based on time to peaks shown in Fig. S1B in (Arkhipov et al. 2018) for the PSC features for connections between different populations.

Excitatory to Excitatory: 5.5 ms
Inhibitory to Excitatory: 8.5 ms
Excitatory to Inhibitory: 2.8 ms
Inhibitory to Inhibitory: 5.8 ms

#### LGN Input

The weights for the 3,506,880 connections from LGN to V1 were tuned based on the target post-synaptic currents from literature (Lien and Scanziani 2013; Ji et al. 2015). The average post-synaptic currents of each population were tuned to match the literature target currents. The small tuning network was created for each population with 100 nodes for each GLIF3 model in the population. As the average rheobase of the GLIF3 models is bigger than the ones from experimental data, we scaled the weights from LGN to V1 by the factor of 1.36, which is average ratio between average rheobases of GLIF3 models and experiment data over all the populations.

In this study, the LGN inputs were generated from stimuli presenting a half second of grey screen and a following 2.5-second drifting grating. The drifting grating is with temporal frequency TF = 2Hz, spatial frequency SF = 0.04 cpd, contrast = 80%, and orientation = 270°. Additional drifting grating stimuli (with a different SF = 0.02 cpd and different contrasts as 5%, 10%, 20%, 40% and 80%) were also used for the multiple neuron perturbations.

#### Background Input

This background (BKG) input represents long range inputs from areas other than LGN. It is to simulate the effect of other parts of the brain on V1, which was in the form of Poisson spike train with firing rate being set as 1,000 Hz. The weights for 230,924 connections from BKG to V1 were initially estimated based on rheobases of the target V1 nodes and then tuned to match the spontaneous firing rate of each population measured from experiments (Durand et al. 2016; Niell and Stryker 2008). Also, the number of synapses of each of such edges were estimated based on the dendritic length of the cell. The tuning of background to V1 weight is to match the spontaneous firing rates from experiments. During the tuning, in views of the too strong background responses from E2/3 and E5 neurons, as well as Pvalb neurons in layers 4, 5 and 6, the weights for the two populations of excitatory neurons, and Pvalb neurons in the three aforementioned layers were scaled by the factors of 0.8 and 0.5. With automated tuning and scaling, the V1 model was tuned to match spontaneous activity during grey screen stimulus from experiment data.

### Cell Type Perturbation

We first performed perturbation simulation experiments at the cell type population level to study the functional roles of different types of inter-laminar interactions. The experiments included population silencing, population activation and local functional population perturbation with current injections to different subsets of population groups. Details of each of these simulation experiments are described as follows.

#### Whole Cell Type Population Perturbation

In the initial perturbations, with negative currents injected to the whole population, one of the 17 populations in the V1 model was silenced. We also examined injecting positive currents to the whole population, with L6 excitatory neurons and L6 Pvalb neurons as a demonstrative example. The strengths of the injection for L6 populations are 0.5 times of the rheobase of the population i.e., E6 or I6Pvalb.

An additional analysis metric we are using for the population silencing is the Optogenetic Modulation Index (OMI). It is computed for every cell and then averaged over the analyzed cells. The OMI of a neuron is defined as:

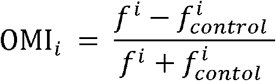

where *f^i^* is the firing rate of neuron *i* in perturbation simulation, and 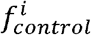 is the one in no-perturbation simulation. Negative OMI indicates suppression of activity due to perturbation (OMI = −1 means that the cell is fully suppressed), OMI = 0 means no change of activity, and positive values indicate elevated activity due to perturbation.

#### Functional Population Perturbation

In the second set of perturbation, subpopulations of excitatory populations in layers 2/3 were perturbed by injecting positive currents three times the rheobase of the population. The perturbed neurons in each simulation were selected with tuning angles in the ranges 270°±45°, 270°±22.5° and 270°±11.25°, and within different radii (i.e., 50μm, 100μm, 150μm, 200μm and 250μm) from the center of the V1 model. The same ten trials of the LGN drifting grating stimuli were run for the simulations.

Firing rate change (Δ*f*) is used to analyze the perturbation results. The response changes were computed for the 17 populations in the model with Δ*f* only computed for those neurons of the same preferring tuning angles and the same radius as the perturbed neurons. The results are shown as heatmaps indicating the relations between Δ*f* and radius (Fig. 2), which are separated by the ranges of tuning angle and perturbed populations (i.e., E2/3). The firing rate changes for all neurons (including the perturbed E2/3 neurons) in the core of the V1 model of each population are shown in Fig. S4. Additional analysis using OMI is shown in Figs. S5 and S6.

### Single Neuron Perturbation

#### Target Neurons

In the single neuron perturbation, positive current was injected to one target Cux2 excitatory neuron in layer 2/3 for each simulation. The target neuron was selected within 50μm radius from the center of the V1 model and as close as the desired preferred angle within 5°. Two desired preferred angles, i.e., 270° and 180°, were used. For each desired preferred angle, 50 different simulations were run, each with a different perturbed neuron. For perturbed target neuron, 10 trials of stimuli were run. The visual stimulus is a drifting grating with TF = 2Hz, SF = 0.04 cpd, contrast = 80%, orientation = 270°. One set of stimulated neurons are aligned to the visual stimulus, while the other is orthogonal. Additional desired preferred tuning angles were also simulated to compare with the observations reported in (Chettih and Harvey 2019), with results shown in Fig. S8. The injecting currents to the target neuron were set as three times of the rheobase of the target neuron models.

#### Neighboring Neurons

To measure the effect of the injection to the target neuron, the activity of neighboring neurons within depth (Y axis) ranges [−50.0 μm, 50.0 μm] around the target neuron were analyzed. The analyzed neighboring neurons were within the 300 μm radius on the horizontal plane. Layer 1 neighboring neurons within this range were not included in the analysis.

#### Grouping Neighbor Neurons

To analyze the effect of single neuron perturbation in different aspects including distance and angle, the neighbor neurons were grouped by the following two criteria:

- **Angle_diff** (in degree): difference of tuning angles between target neuron *θ_target_* and neighbor neurons *θ_neighbor_*.
- **2D Distance** (in μm): distance from the neighbor neuron to target neuron projected on the horizontal plane.

#### Effect Measurements

To measure the effect of the perturbation on the target neuron, two metrics were used:

- **Firing Rate Change** (Δ*f* in Hz): firing rate change of a neighboring neuron with current injection to the target neuron and without current injection to the target neuron, i.e., 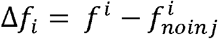 for the same LGN stimulus and BKG input
- Δ**Activity**: the activity change metric (Chettih and Harvey 2019) for neuron *i* in the trial *j* of the simulation is:

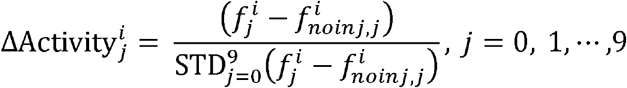

where the firing change 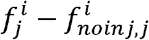 of the *j*th trial is divided by the standard deviation 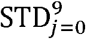 of such firing rate changes over the 10-trial simulations for the same LGN stimulus. Note that, different from the metric in (Chettih and Harvey 2019) using average activity of control sites as the control, the fully-control simulated V1 enable us to get the exact activity of each neighbor neuron for each trial without perturbation and get the exact difference between the perturbed activity and no-perturbed activity.

#### Effect Summation

The evaluation metrics introduced above were averaged across 10 trials for each neighbor neuron. The averaged metric values of all the neighbor neurons were binned into different groups. The number of the bins for distance-based analysis is 30, while the angle-based one is 4. The average metric value for each group of the neighboring neurons (i.e., average value of each bin) is used for evaluation the effect of the perturbation on the target neuron to the neighbor neurons of the same group. Also, for firing rate change metric, excitatory and inhibitory neighbor neurons were separated in the analysis, with results demonstrated in Figs. 3D, 3E, S7C and S7D. To compare with results reported in (Chettih and Harvey 2019), excitatory and inhibitory neighbor neurons were combined together and analyzed using the metric ΔActivity, with results shown in Figs. 3C and S7B. The target neuron in each simulation was excluded in the activity change analysis.

### Multiple Neuron Perturbation

Multiple neuron perturbation simulations were conducted to explore how activity change of one or more cortical neurons could influence nearby cortical neurons and network activity. We targeted excitatory neurons in layer 2/3 and analyzed the activity change of the surrounding neighboring neurons with close and similar prefer tuning angles under perturbation on the target neurons.

#### Target Neurons

Positive current with strength as 3 times of the rheobase was injected to one or more target Cux2 excitatory neurons in layer 2/3. The target neurons were selected within 200 μm radius around the center of the V1 model and the desired preferred angles within 11.25° around 270° which is the input stimulus direction. The numbers of perturbed neurons are 1, 2, 3, 5, 10, 20, 30, 50 and 100. For each number of perturbed neurons, ten different sets of the target Cux2 neurons were randomly selected. For each set of target neurons, 10 trials with different instantiations of the random inputs were run. The visual stimulus is a drifting grating with TF = 2Hz, SF = 0.04 cpd, contrast = 80%, orientation = 270°. Additional simulations were also conducted with SF = 0.02 cpd and five different contrasts as 5%, 10%, 20%, 40% and 80%. Note that the radius and angle range for the selection of target neurons were chosen to ensure enough neurons for perturbation simulations.

#### Neighbor Neurons

To measure the effect of the perturbation, the activities of neighboring neurons of each cell type population were analyzed. The analyzed neighboring neurons were with the 200 μm radius on the horizontal plane. The neighboring neurons within the following three different ranges of preferred tuning angles were analyzed and compared.

- 270°±45°
- 270°±11.25°
- 270°±45°, excluding 270°±11.25°

We also analyzed neighboring neurons within 16 different ranges of preferred tuning angles around the whole circle with range bins being 22.5° and center being at 0°, 22.5°, 45° etc (Fig. S10).

#### Effect Measurements and Summation

Firing Rate Change (Δ*f* in Hz) was used and it was averaged across 10 trials. The averaged values of all the selected neighbor neurons of a cell type were averaged again to get the average Δ*f* for that cell type population. The average population Δ*f* was then averaged across the 10 randomized selections of target neurons. The overall Δ*f* was compared across different numbers of target neuron simulations as shown in Fig. 4 in the forms of heatmaps for all 17 populations in the V1 model and curve plots for the E2/3 population of different ranges of preferred angles. The stimulated target neurons were excluded from population averages.

## Acknowledgments

We wish to thank the Allen Institute for Brain Science founder, Paul G. Allen for his vision, encouragement and support.

## Supplemental Information

### Supplemental discussion on the similarities and differences between simulated and observed single neuron perturbations

An additional analysis (Figs. S7-S9) was conducted for the single E2/3 neuron perturbation simulations, to further compare the simulation results with experiment results reported in (Chettih and Harvey 2019). The comparative analysis shows consistency between simulations and experiments in term of general suppression, higher order terms in distance dependence (Fig. S8) and the dependence of interactions on the orientation difference between recorded neuron’s preferred tuning and visual stimulus (Fig. 3E) when the stimulated neuron aligns with the visual stimulus. That is, neurons preferentially responding to the visual stimulus are most suppressed as shown in Figs. 3E and discussed in the Results section. There are differences in higher order effect between stimulated and recorded neurons when the stimulated and recorded neurons and visual stimuli are at different angles.

However, when it comes to higher-order observation regarding the interaction between the direction of recorded neuron and the preferred direction of the stimulated neuron, our V1 model illustrated distinct result with the experiment in (Chettih and Harvey 2019). The experiment results showed the decrease in gain described in the 1st-order suppressive effect was greatest when the tuning preference of stimulated neuron matched the presented visual stimulus, while our model simulation results did not show such higher-order effect, but the opposite is shown in Fig. S9. A potential reason for the difference of the higher-order effect between the simulation and experiment is the complexity of the neuron preferences. Sparse coding (Olshausen and Field 1996a, 1997) predicts a like-to-like inhibition in absence of noise and spikes. If the visual code representing a feature does not map directly to the activity of one neuron, but an average over a subpopulation of neurons with identical codes, one would expect a stimulation of one neuron to not have an inhibitory effect on identically tuned neurons. Such an effect is observed both in simulations (Fig. S10 A) and in experiments and can be described as a “Mexican hat” for the interactions in functional space. In our model, the tuning properties of individual neurons are probably not as varied as the ones in real mouse V1. Our V1 model has been thoroughly tested to match the biological observations of representation of different directions of movements. It is possible that on other features, for which we did not have a diversity of stimuli presented, the preferences of cells are too similar for them to engage in meaningful competition. As such, if the functional space for biology has higher dimensionality than the functional space in the simulations, the number of neurons with similar but not closely-aligned tuning will be much higher in experiments, and will be most often seen.

### Supplemental figures

**Figure S1.**
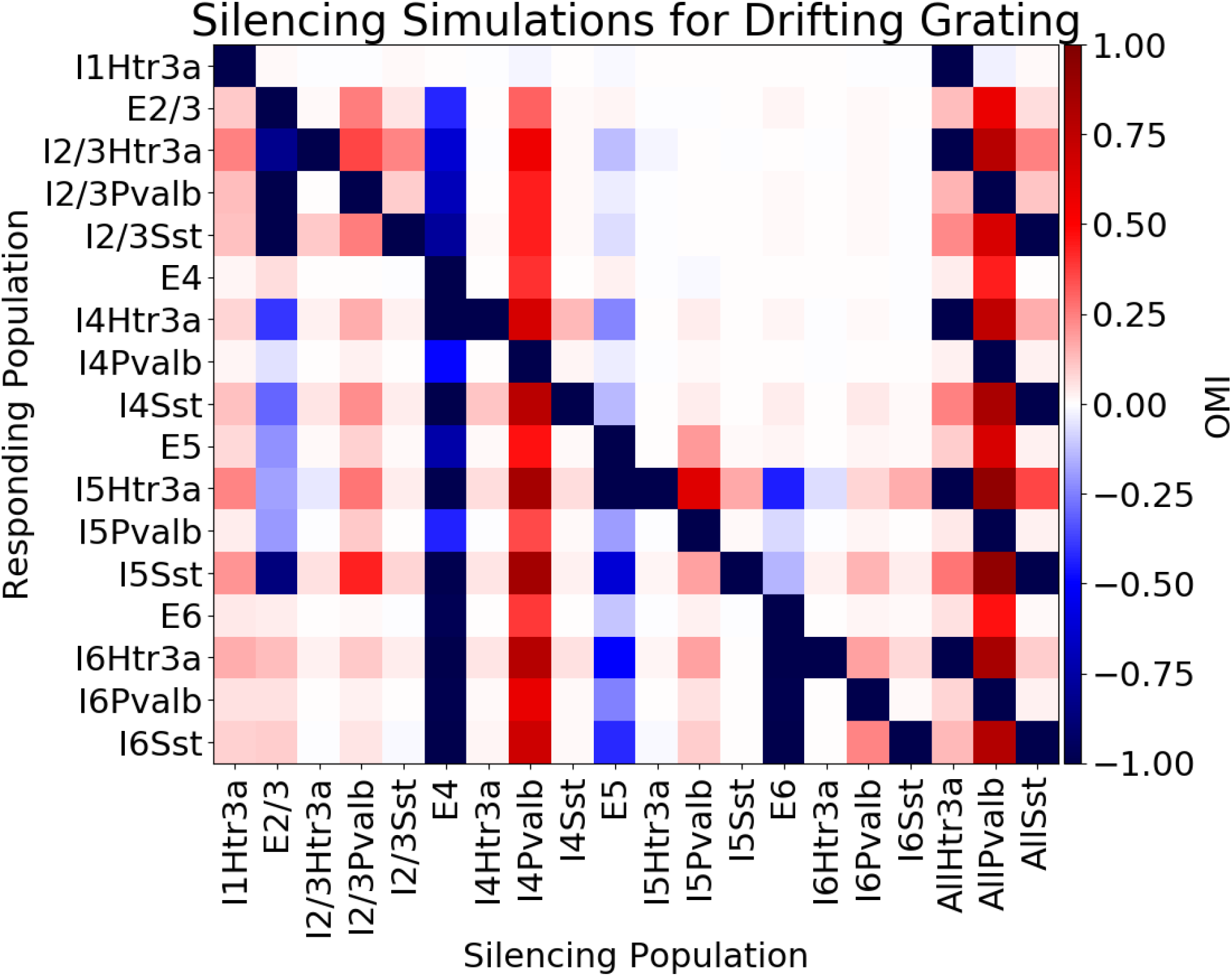
Summary of silencing simulations during presentation of 0.5 s of grey screen and 2.5 s of a drifting grating (TF = 2Hz, SF = 0.04 cpd, contrast = 80%, orientation = 270°). Labels along the horizontal axis indicate the silenced populations. Labels along the vertical axis indicate the populations for which OMI is computed. The entries on the bottom right (i.e., “allHtr3a” “allPvalb” and “allSst”) refer to perturbations where multiple populations were silenced together (e.g., “allSst” means silencing Sst neurons in all layers). The OMIs showing in the heatmap are the average values across stimulus trials and neurons in a population. Silencing E4 neurons leads to suppression of activity throughout the layers, whereas silencing other excitatory neurons disinhibits excitatory neurons in other layers. Furthermore, silencing inhibitory populations leads to elevated activity across the column in most cases.

**Figure S2.**
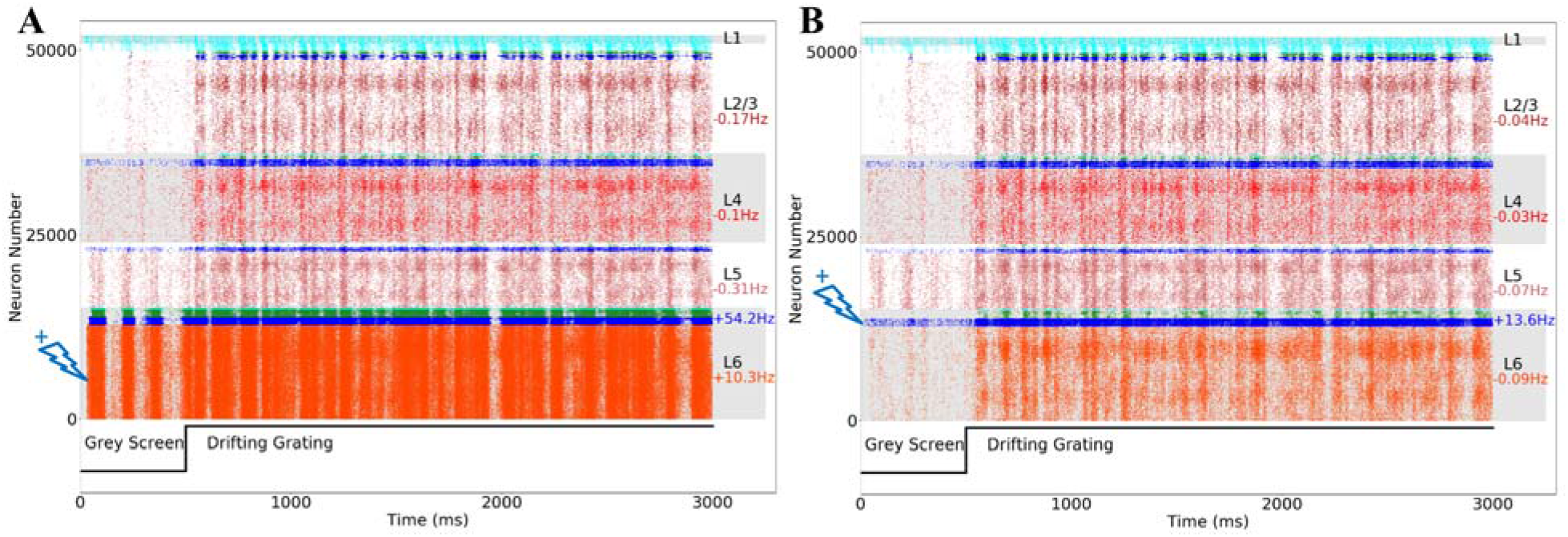
Raster plot of whole population perturbations of neurons on layer 6 following presentation of a drifting grating. (**A**) Raster plot of activation of E6 with half of the rheobase of the target population, resulting in excitation of layer 6 Pvalb and suppression of other upper layer excitatory cells. (**B**) Raster plot of activation of I6Pvalb neurons with half of the rheobase of the target population, suppresses all excitatory cells across the column.

**Figure S3.**
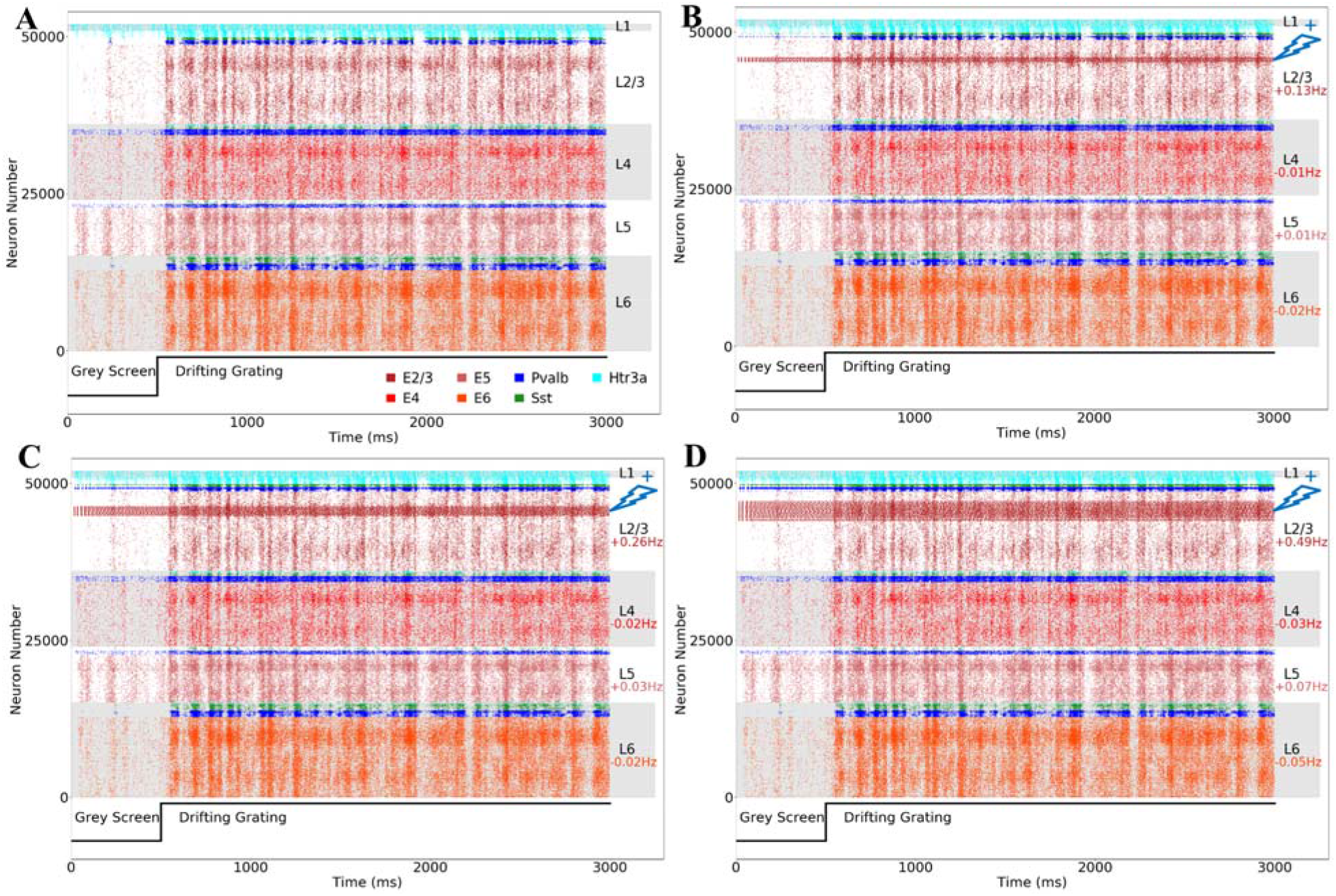
Raster plot of population perturbations of E2/3 during presentation of a drifting grating, with the injected current being 3 times of the rheobase of the target population. (**A**) Baseline. (**B**) The perturbations applied to neurons that prefer motion in three directions – 270°+/−11.25° (**C**), 270°+/−22.5° or (**D**), 270°+/−45° – and situated within 100 μm of the center. As more neurons are perturbed, the stripe of activated increases becomes wider.

**Figure S4.**
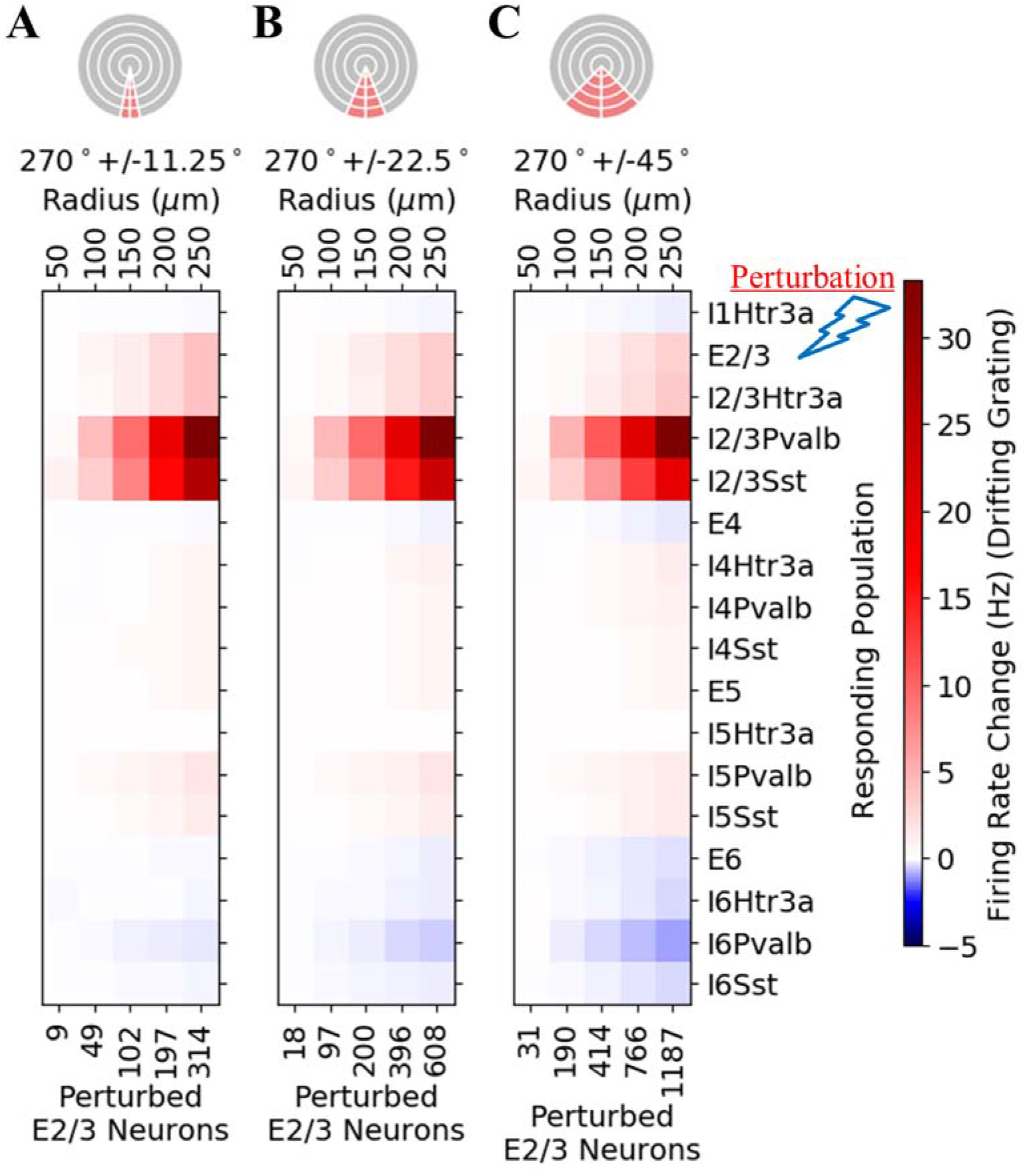
The exact same perturbation as in Fig. 2 with the sole difference that is Δ*f* averaged over all orientations within the radii indicated rather than for the different functionally defined populations. The activation of a functionally co-tuned subpopulation of E2/3 neurons lead to a robust activation of inhibitory neurons in the same layer. They produce an inhibition of dissimilaryly tuned excitatory neurons, such that at the population level, the overall effects are small.

**Figure S5.**
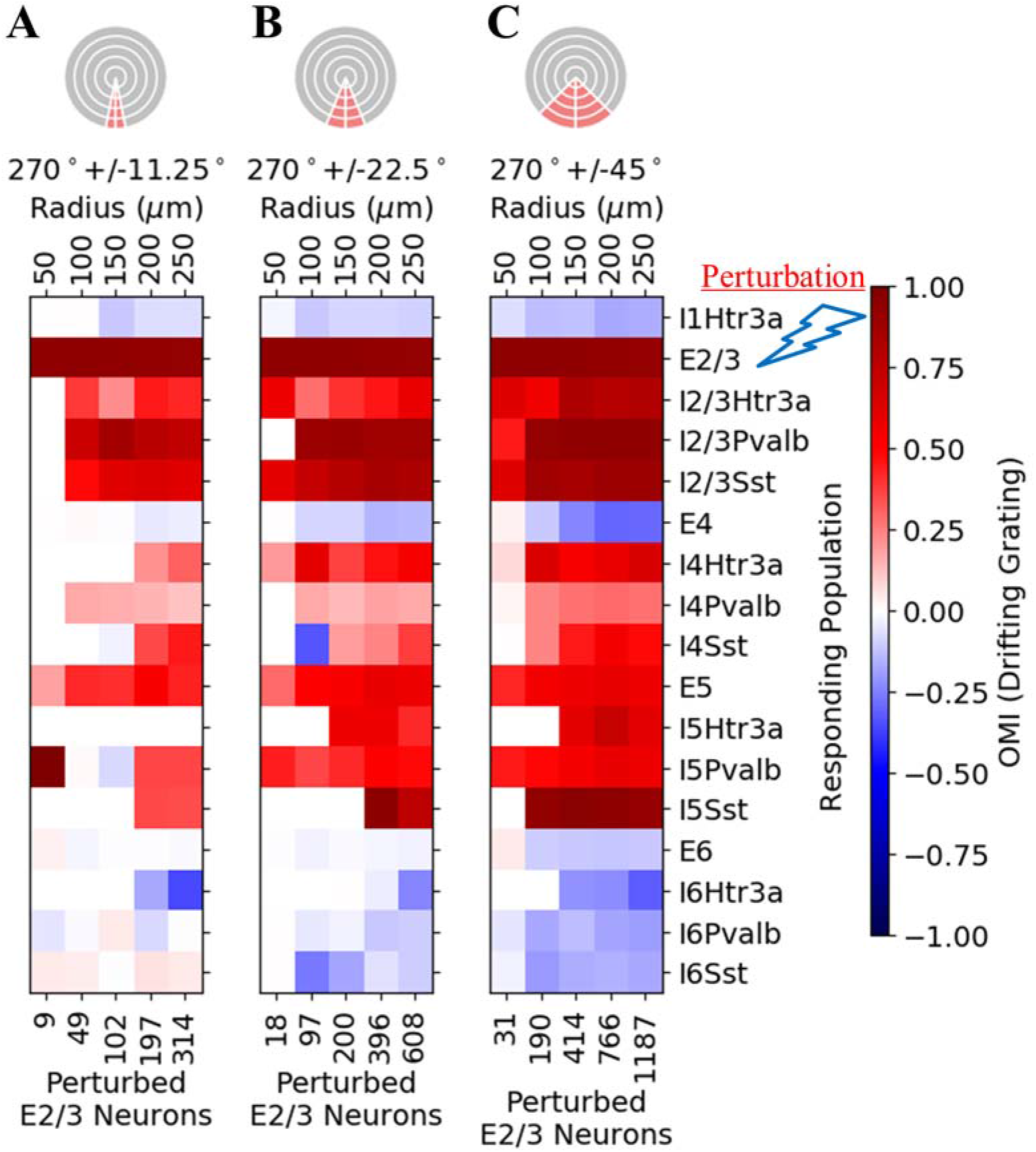
The same perturbation as in Fig. 2, S2 and S4 but using OMI as the analysis measurement rather than Δ*f*. The OMI for the different genetically defined populations is evaluated over the neurons that have the same direction preference and radial positions as the perturbed populations. The figure indicates successful imprinting by exciting E2/3, leading to remarkably distinct patterns of activation. The perturbation effect on the whole column becomes stronger as the number of perturbed neurons increases

**Figure S6.**
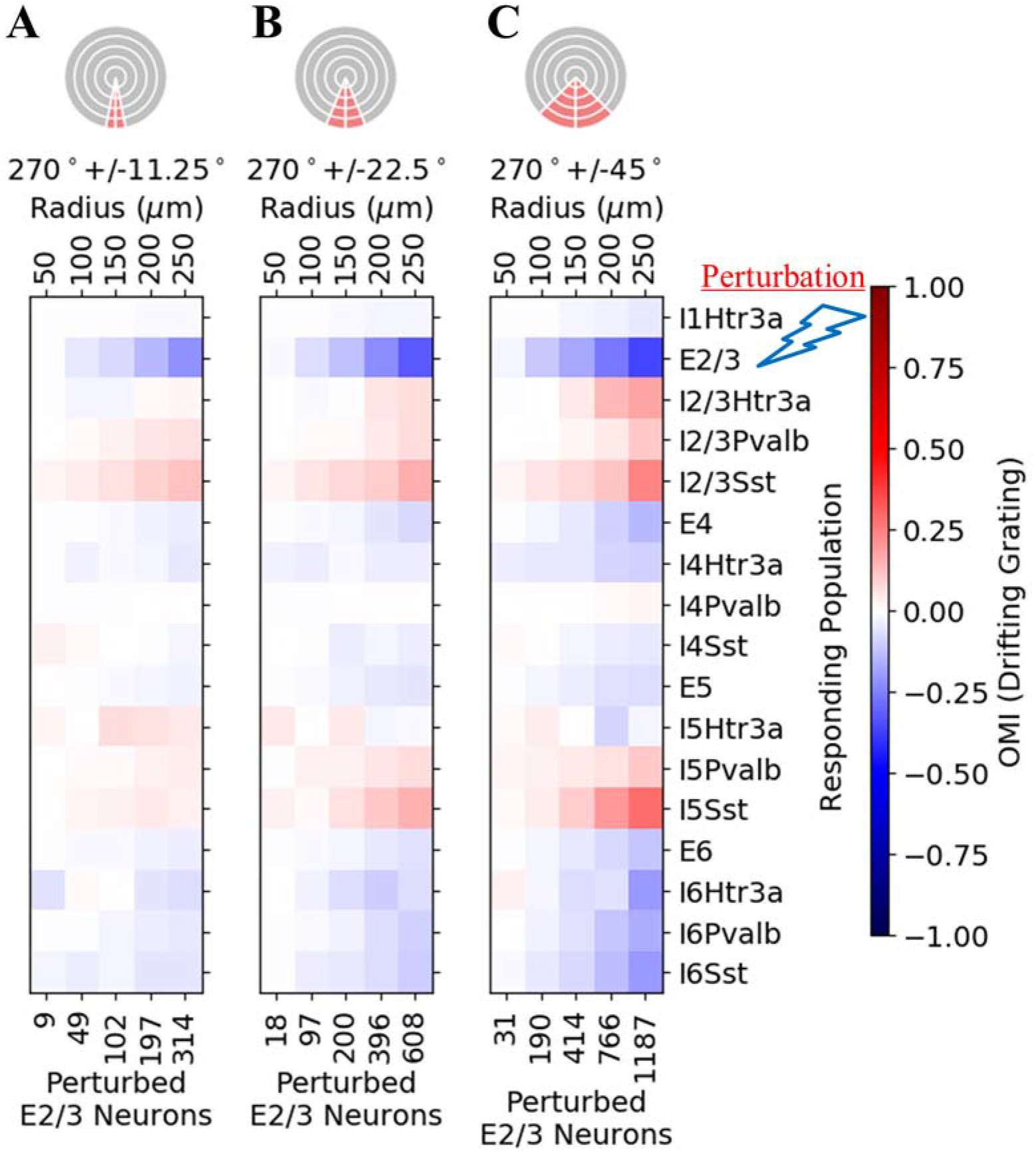
The same perturbation as in Fig. 2, S2-S4, but using OMI evaluated for all neurons within the core (unlike Fig. S5 in which the computation is restricted to functionally defined sub-populations).

**Figure S7.**
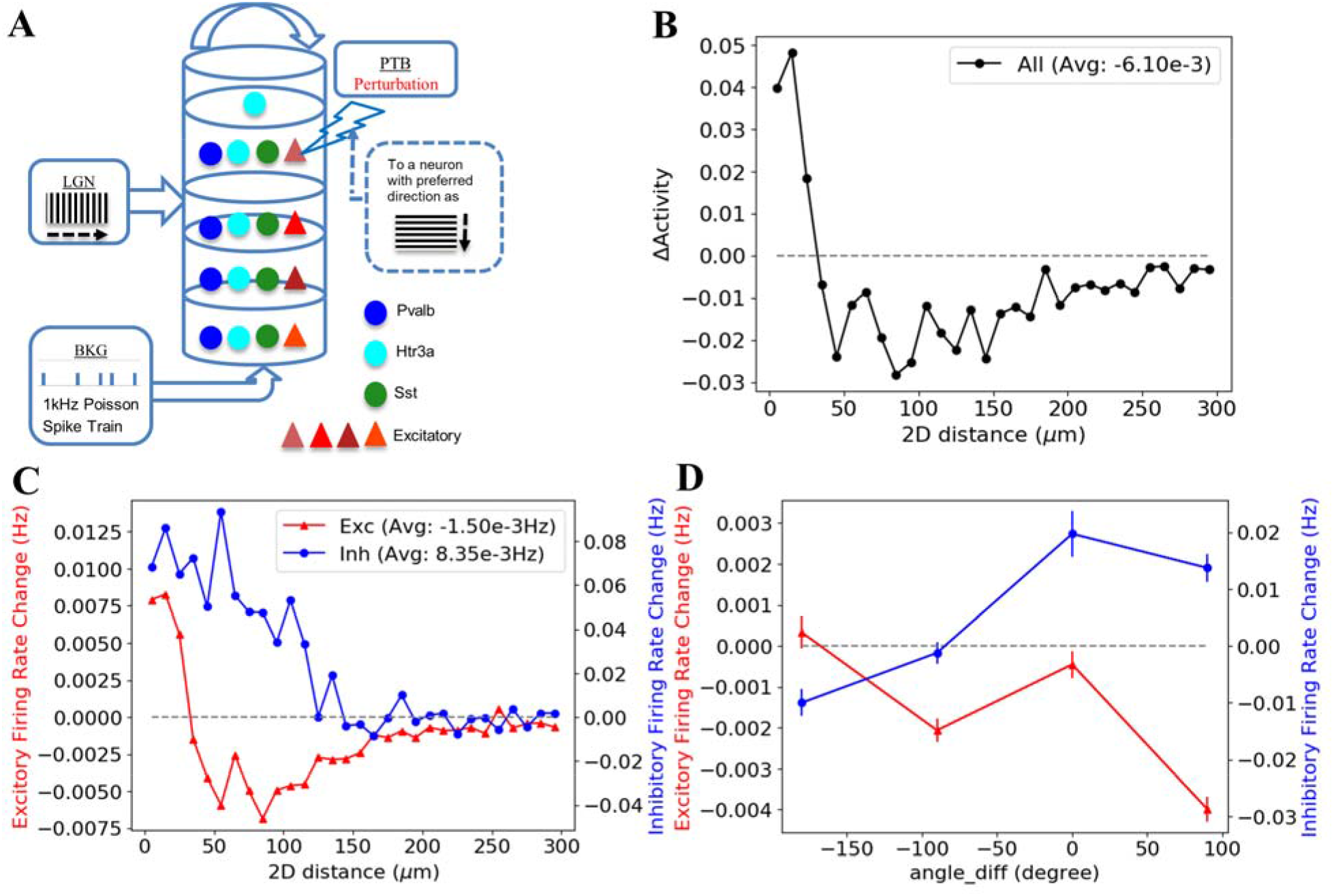
Using the V1 model to study effects of single-neuron perturbations on the population activity by stimulating an excitatory neuron with preferred direction orthogonal to the direction of the input stimulus (instead of same preferred direction in Fig. 3). This leads to similar patterns of distance dependence (**B**, **C**) to Figs. 3C and 3D, but substantial differences in orientation dependence: excitatory neurons tuned for the stimulus (but not the preferred angle of the target neuron, i.e., angle_diff = 90°) were suppressed more than those dissimilarly tuned (**D**).

**Figure S8.**
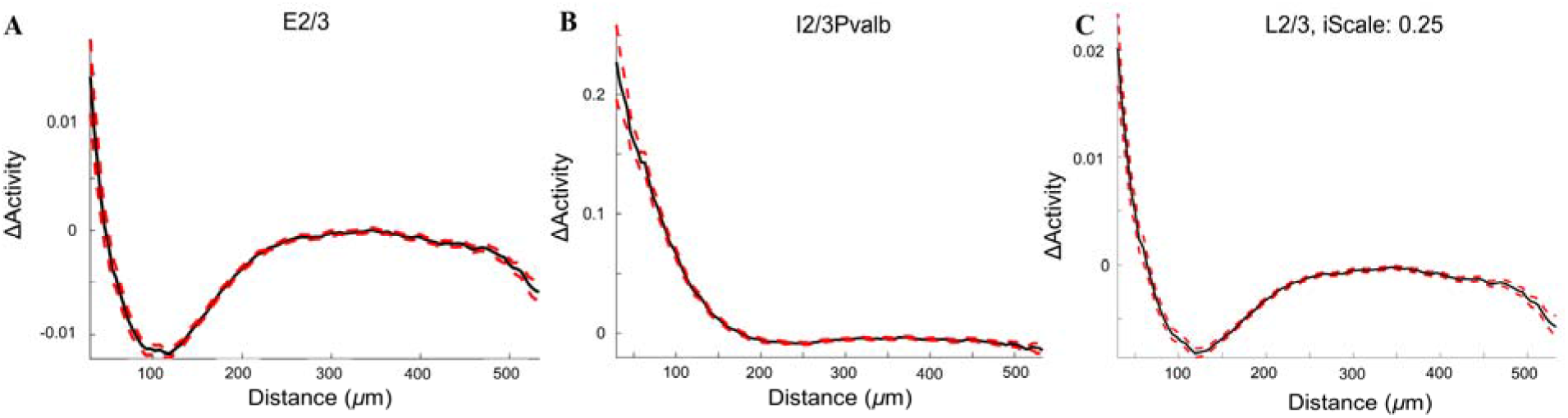
Distance dependent activity change of single neuron perturbation using the V1 model. The activity change measure is defined as firing rate changes normalized by standard deviation of firing rate changes of the 10 trials of each of the drifting grating stimuli simulations, which is similar to ΔActivity used in (Chettih and Harvey 2019). The measure plot is depending on distance from stimulated neuron. (**A**) Activity change of excitatory neurons in layer 2/3 with standard error shown as red dashed lines. We observe the center-surround effect similar to the ones shown in Figs. 3D and S7C, and is consistent with the findings in the experiment (Chettih and Harvey 2019). (**B**) Activity change of inhibitory neurons with Pvalb neurons in layer 2/3 shown as an illustrative example. We again observe the center-surround effect similar to the ones shown in Figs. 3D and S7C, and is consistent with the findings in the experiment (Chettih and Harvey 2019) as well. (**C**) Activity change of all excitatory and inhibitory neurons in layer 2/3. We observe the center-surround effect similar to the ones shown in Figs. 3C and S7B and is consistent with the findings in the experiment (Chettih and Harvey 2019). From the figure, we can see the E-I crossover is at 70μm which is consistent with the one reported in the experiment (Chettih and Harvey 2019). Note that measurements from inhibitory neurons were scaled down by a factor of 0.25 to match the experiment setting as some fraction of inhibitory neurons were included in the experiment data collected based calcium signals from excitatory neurons.

**Figure S9.**
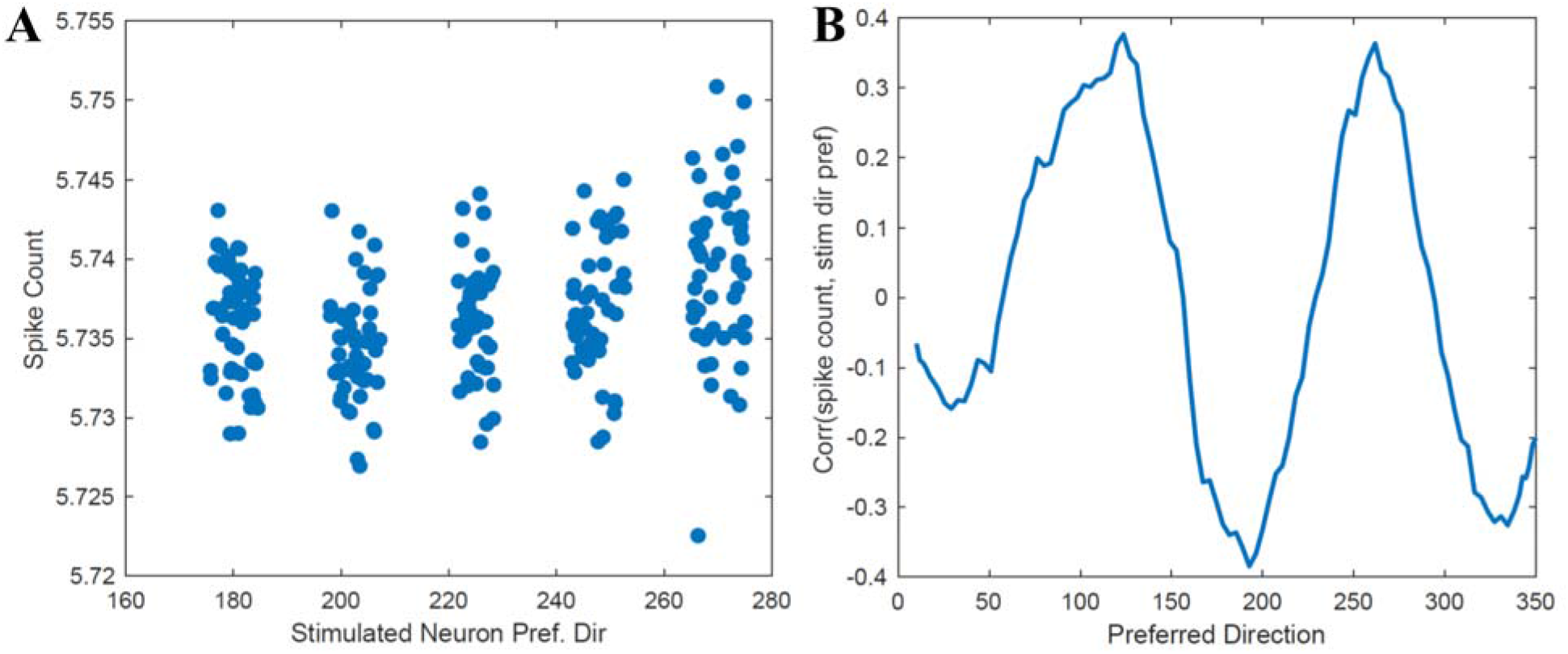
Tuning angle dependency analysis for single neuron perturbations using the V1 model for simulations of 5 sets of stimulated neurons with preferred orientations as 180°, 202.5°, 225°, 247.5° and 270°. (**A**) Spike counts of responding neurons with preferred tuning angle near 270° (i.e., the orientation of the visual stimulus) with respect to preferred orientation of every stimulated neuron. We can observe variability between neurons with nearby preferences, but on top of this variability it is very clear that there are more spikes in these neurons when a 270° preferring neuron was stimulated. This suggests that the stimulus gain is greatest when stimulated neuron preference and presented visual stimulus coincide. This is counter to the strongest suppression observation in the experiments as shown in Fig. 4e in (Chettih and Harvey 2019). This relation is summarized into one number, the correlation between the spike count of the neurons preferring 270° and the stimulated neuron preferred direction. (**B**) Correlations of the spike counts of neurons as a particular preferred direction with the preferred direction of the stimulated neuron (the correlation for the plot in A is represented as the point in B at 270°). The figure shows that neurons around 90° and 270° (i.e. those with orientation preference matching the visual stimulus) have greater responses as the stimulated neuron’s presence goes from 180° to 270°, and the opposite is true for neurons at 0° and 180°. This again shows the opposite 3^rd^-order effect reported in (Chettih and Harvey 2019).

**Figure S10.**
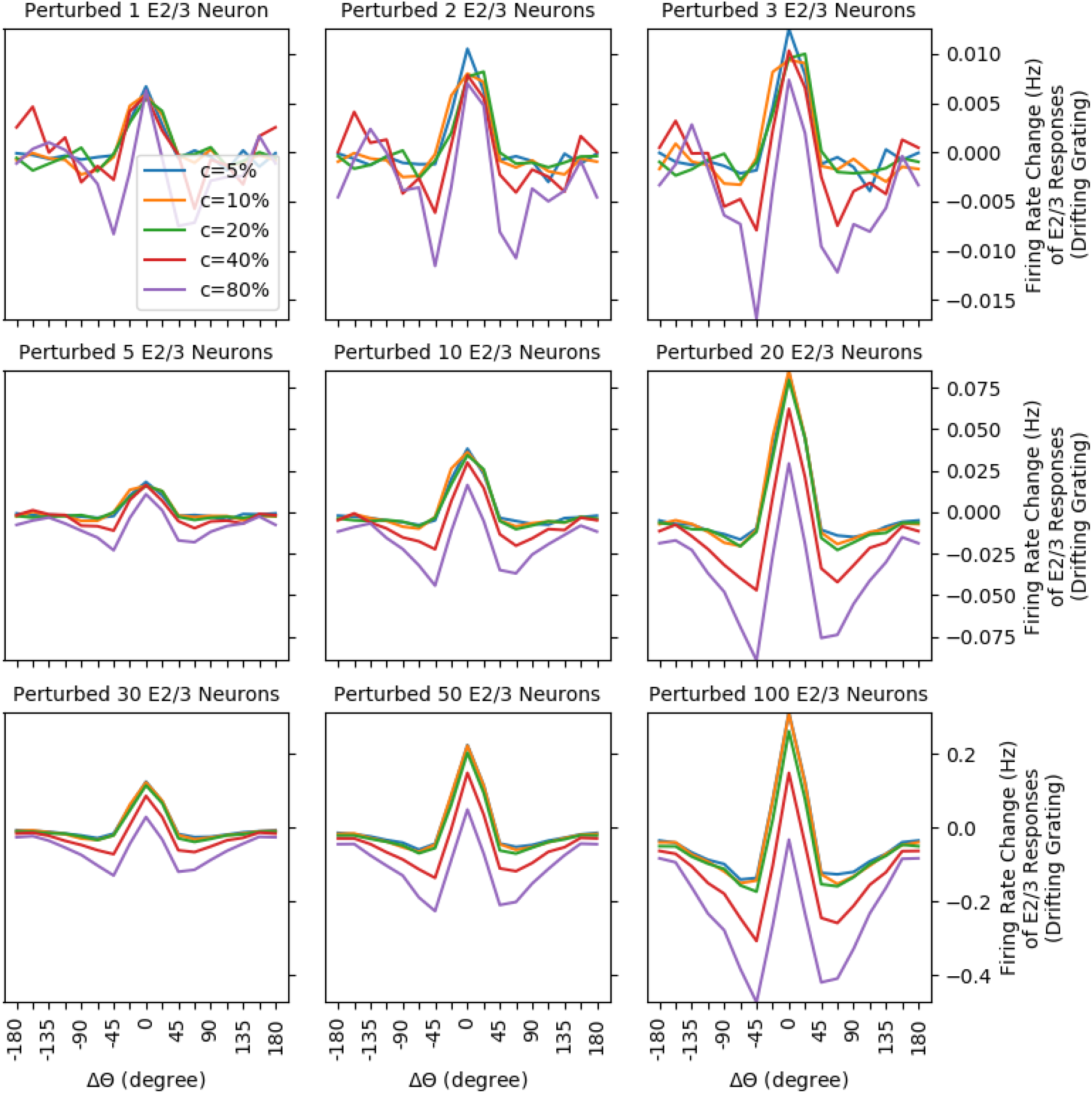
Perturbations of multiple excitatory neurons (from 1 neuron to 100) in layer 2/3 during drifting grating stimulation. The perturbed neurons were selected such that their preferred direction closely matched the direction of the drifting grating: 270°+/−11.25° and within 200 μm radius. The plots show activity changes of E2/3 neurons dependent on the angle difference between the analysis of neurons and the stimulus direction (i.e., 270°). Each of these plots is for simulation of a particular number of perturbed neurons. In the plot, labels along x axis indicate such angle differences in 22.5° bins, and labels along the y axis indicate the firing rate changes of the analyzed neurons. The curves were color coded for five different contrasts (5%, 10%, 20%, 40% and 80%) for drifting grating with 0.02 cpd spatial frequency. We can see Mexican hat shape curve for activity changes of E2/3 neurons centering around the visual stimulus direction. Closely aligned E2/3 neurons (within 11.25°) were the ones most activated, while the ones similarly tuned (within 45°) show mostly-decreasing activities. Such an effect was decreased as the contrast increased and also as the number of perturbed neurons decreased.

